# Single-dose immunisation with a multimerised SARS-CoV-2 receptor binding domain (RBD) induces an enhanced and protective response in mice

**DOI:** 10.1101/2021.05.18.444622

**Authors:** Ralf Salzer, Jordan J. Clark, Marina Vaysburd, Veronica T. Chang, Anna Albecka, Leo Kiss, Parul Sharma, Andres Gonzalez Llamazares, Anja Kipar, Julian A. Hiscox, Andrew Owen, A. Radu Aricescu, James P. Stewart, Leo C. James, Jan Löwe

**Affiliations:** MRC Laboratory of Molecular Biology, Cambridge, UK; Institute of Infection, Veterinary and Ecological Sciences, University of Liverpool, Liverpool, UK; Laboratory for Animal Model Pathology, Institute of Veterinary Pathology, Vetsuisse Faculty, University of Zurich, Zurich, Switzerland; Department of Pharmacology and Therapeutics, Centre of Excellence in Long-acting Therapeutics (CELT), University of Liverpool, UK

**Keywords:** COVID-19, SARS-CoV-2, coronavirus, vaccine candidate, subunit vaccine, RBD, Dps

## Abstract

The COVID-19 pandemic, caused by the SARS-CoV-2 coronavirus, has triggered a worldwide health emergency. So far, several different types of vaccines have shown strong efficacy. However, both the emergence of new SARS-CoV-2 variants and the need to vaccinate a large fraction of the world’s population necessitate the development of alternative vaccines, especially those that are simple and easy to store, transport and administer. Here, we showed that ferritin-like Dps protein from hyperthermophilic *Sulfolobus islandicus* can be covalently coupled with different SARS-CoV-2 antigens via the SpyCatcher system, to form extremely stable and defined multivalent dodecameric vaccine nanoparticles that remain intact even after lyophilisation. Immunisation experiments in mice demonstrated that the SARS-CoV-2 receptor binding domain (RBD) coupled to Dps (RBD-S-Dps) shows particular promise as it elicited a higher antibody titre and an enhanced neutralising antibody response compared to the monomeric RBD. Furthermore, we showed that a single immunisation with the multivalent RBD-S-Dps completely protected hACE2-expressing mice from serious illness and led to efficient viral clearance from the lungs upon SARS-CoV-2 infection. Our data highlight that multimerised SARS-CoV-2 subunit vaccines are a highly efficacious modality, particularly when combined with an ultra-stable scaffold.

## INTRODUCTION

On 11 March 2020 the World Health Organisation declared the COVID-19 outbreak, caused by the SARS-CoV-2 virus, a pandemic (Cucinotta and Vanelli, 2020). Since then, COVID-19 and the efforts to contain it have changed the lives of unprecedented numbers of people. For example, in April 2020 3.9 billion people were affected by lockdown measures aimed to cut or at least reduce the chain of transmission with widespread negative impacts on employment, education and other health issues. According to the Johns Hopkins University there have so far been 151M confirmed COVID-19 cases globally (May 2021) and virtually every country has been affected. Officially, 3.2M people have died from SARS-CoV-2 infection (2021a, 2021b).

SARS-CoV-2 belongs to the family of *Coronaviridae*, which contain a positive-stranded RNA genome (Pal et al., 2020). The RNA is enveloped by a membrane that harbours four coat proteins (Fig. 1A). On the inside of the virus, the nucleocapsid protein (NP) is crucial for RNA packaging and viral release from host cells (Zeng et al., 2020). The Spike protein, which is embedded in the virus’ membranous envelope, is essential for the interaction with human angiotensin-converting enzyme 2 (hACE2) (Ke et al., 2020). It is the interaction with hACE2 that is thought to initiate the process that leads to cell entry of viral RNA and infection (Shang et al., 2020). The Spike protein is translated as a single polypeptide that is proteolytically processed into its two subunits, S1 and S2. The Spike of SARS-CoV-2 is a trimer consisting of three S1-S2 heterodimers (Huang et al., 2020). For membrane fusion between the cell and the virus to occur, two cleavage events within the Spike complex are required (Ke et al., 2020). A protease cleavage site located between S1 and S2 is cleaved by the producer cell’s proprotein convertase furin during virus assembly (Papa et al., 2021) (Fig. 1A). The second cleavage site is located in the S2 domain at position R797, and its hydrolysis by the target cell’s surface protease TMPRSS2 triggers membrane fusion and cell entry (Papa et al., 2021).

**Figure 1.**
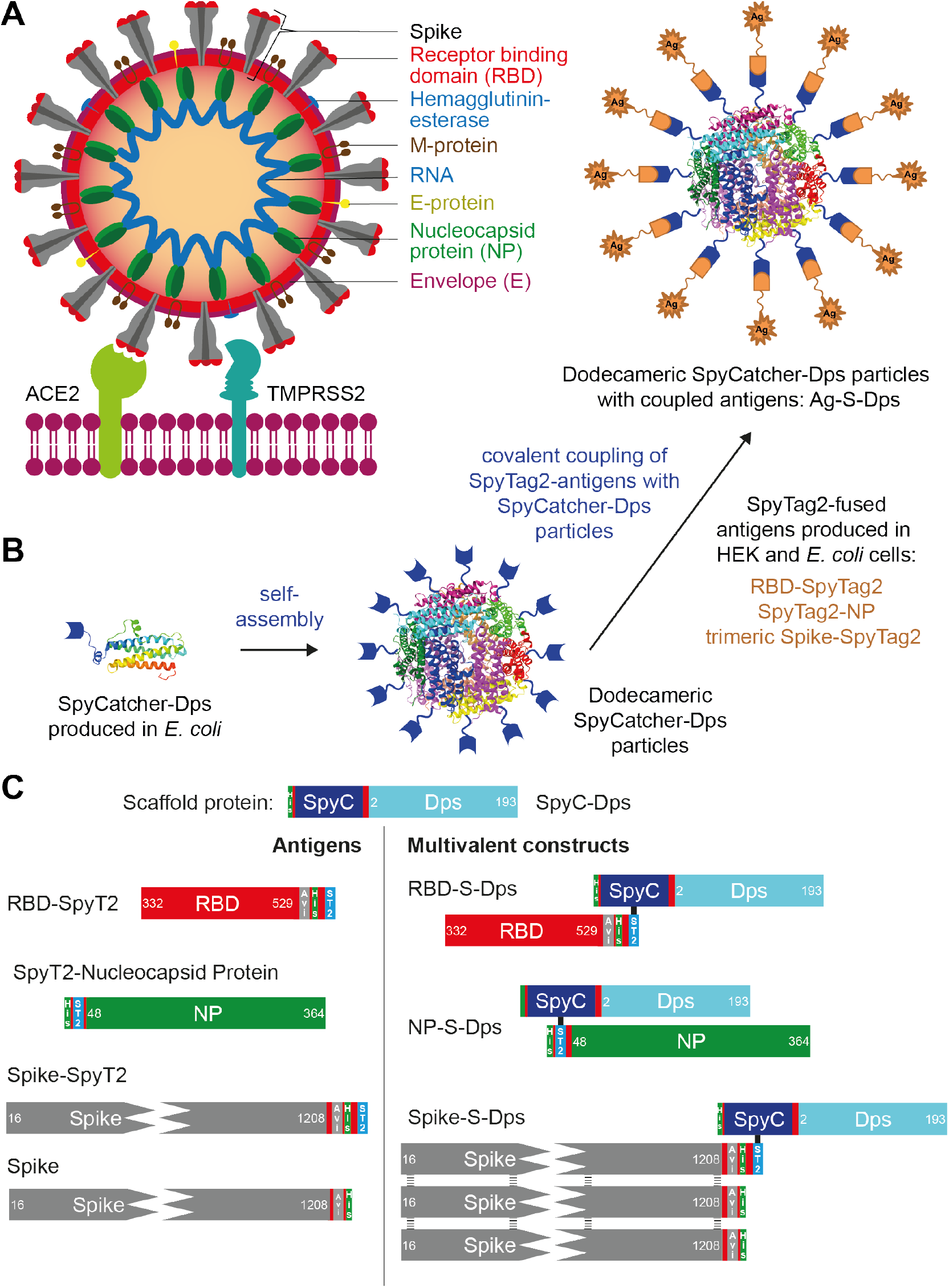
Overview of the multimerisation strategy employed and the antigens and scaffold used. **A**) Cartoon representation of SARS-CoV-2 binding to a human cell membrane. **B**) Schematic diagram of the *Sulfolobus islandicus* Dps and SpyCatcher-based display and multimerisation strategy employed in this study. **C**) Diagram of the proteins used in this work. SpyC is the ΔN1-SpyCatcher domain and SpyT2 is the peptidic SpyTag2 that becomes covalently linked to SpyC upon simple mixing. Stabilised, trimeric Spike/Spike-SpyT2 contained on average only one SpyT2 tag in order to avoid uncontrolled oligomerisation when coupled to Dps.

The SARS-CoV-2 receptor-binding domain (RBD) is located within the S1 subunit of the Spike. It is the RBD that interacts directly with the host cell via the hACE2 receptor (Ke et al., 2020). It is therefore not surprising that antibodies directed against the RBD or overlap with the ACE2 binding region are strongly neutralising, making the RBD a promising subunit vaccine candidate (Ke et al., 2020; Seydoux et al., 2020). The RBD is glycosylated and contains four disulphide bridges that contribute to its stability, necessitating its expression in mammalian cells, as is also the case for the Spike.

To end the pandemic, vaccines are by far the most promising approach and vaccine developments, clinical trials, approvals and mass roll-outs are in progress. So far, until May 2021, 89 COVID-19 vaccines have been tested in clinical trials. Of those, 36 are undergoing safety trials, 27 are in the phase of large-scale testing, 6 vaccines are authorised for limited use, and 8 vaccines are fully approved (2021a). All approved vaccines show good-to-excellent protection against severe illness and preliminary data have shown that virus transmission is significantly reduced in vaccinated individuals (Mahase, 2020; Thompson et al., 2021). Most of the approved vaccines and those in late-stage trials are mRNA-based, vector-based, inactivated viruses or DNA vaccines (Mahase, 2020). Vector- and RNA-based vaccines can often be rapidly developed because they deliver the immunogen coding sequence rather than the immunogen itself. Currently, only one vaccine candidate in late phase trials is a protein-based subunit vaccine, Novavax (Mahase, 2021). Some subunit vaccines are amenable to processes such as lyophilisation that remove the need for a complex storage and distribution cold-chain. As such, they provide substantial advantages over nucleic-acid based vaccines in the quest for complete and global vaccination. A second challenge facing global vaccination is the emergence of viral variants, some of which are more infectious and/or cause more severe illnesses, and reduce the efficacy of existing vaccines (Davies et al., 2021; Ferreira et al., 2021; Kupferschmidt, 2021; Zhang et al., 2021). Repeat vaccinations directed against these variants, but that use the same type of vaccine, could be problematic. This is because immunity is generated against the vaccine vector itself, neutralising it before it can deliver its immunogen cargo (Bottermann et al., 2018). It is anticipated that in future, several different types of vaccines will be required to cope with emerging variants of SARS-CoV-2.

Previous work has shown that protein-based subunit vaccines directed against SARS-CoV-2 deliver high antibody responses in animal models (Tan et al., 2021; Wang et al., 2021). Furthermore, subunit antigens have the potential to deliver a cheaper, boostable and more robust alternative to nucleic-acid based vaccines (Dalvie et al., 2021; Gu et al., 2021; He et al., 2021; Joyce et al., 2021; Kalathiya et al., 2021; Koenig et al., 2021; Ma et al., 2020; Powell et al., 2021; Xiang et al., 2020). To explore the development of stable and efficient subunit vaccine candidates, we covalently linked SARS-CoV-2 proteins expressed in mammalian and bacterial cells with bacterially-expressed Dps from the hyperthermophilic archaeon *Sulfolubus islandicus* (Gauss et al., 2006). Immunisation using SARS-CoV-2 RBD linked to Dps (RBD-S-Dps) proved to be highly effective in eliciting an immune response and to produce neutralising antibodies that inhibit cell entry *in vitro*. Furthermore, transgenic K18-hACE2 mice infected with SARS-CoV-2 were completely protected from serious illness following a single immunisation with RBD-S-Dps.

## RESULTS

### Three multimerised SARS-CoV-2 antigen complexes

We aimed to find a stable, convenient and non-bacterial display scaffold that would allow the display and multimerisation of a range of SARS-CoV-2 antigens (Fig. 1A). Multimerisation has been used for many years to increase the immunogenicity of different antigens through multivalency, and this approach has also been recently shown to work well with SARS-CoV-2 antigens (Dalvie et al., 2021; Kalathiya et al., 2021; Powell et al., 2021; Wang et al., 2021).

For the purpose of stable multimerisation, we identified Dps (ORF SIL_0492) from *Sulfolobus islandicus*. The source organism is an archaeon, which prefers pH ∼3 and, as a hyperthermophile, has adapted to grow optimally at temperatures of around 80 °C. The intrinsic thermostability and environmental robustness of *S. islandicus* Dps make it an outstanding candidate for the development of a multimerisation scaffold. Dps, a member of the ferritin-like protein family, self-assembles into hollow, dodecameric spheres with 12 subunits, which are roughly 10 nm across (Gauss et al., 2006). Most ferritins assemble larger spheres with 24 subunits. Also, in contrast to *bona fide* ferritin scaffolds, both the N- and the C-termini of Dps are accessible on the outside of the sphere.

We aimed to test whether Dps could efficiently display Spike, RBD and also NP antigens of SARS-CoV-2 (Fig. 1A). Spike and RBD cannot be expressed in folded form in *E. coli*, whereas NP as well as Dps express and fold well in *E. coli*. Expression of soluble and multimeric antigens genetically fused to Dps in mammalian cells (or *E. coli*) was unsuccessful, therefore we decided to employ the SpyCatcher/SpyTag system to attach Dps to different antigens. The SpyCatcher/SpyTag system forms isopeptide bonds between amino acid side chains of the catcher domain and the peptidic tag (Brune and Howarth, 2018; Zakeri et al., 2012). ΔN1-SpyCatcher (Bruun et al., 2018) was fused genetically to the N-terminus of Dps, separated by an eight amino acid long GS linker and a hexa-histidine tag added for purification purposes (SpyC-Dps, Fig. 1B & C). We chose N-terminal linkage to Dps, SpyC-Dps, rather than Dps-SpyC since the coupling reactions were more efficient, but we did not explore this in any detail. Both the N- and C-terminus of Dps are on the outside of the sphere and are accessible for covalent coupling. For the antigens, SpyTag2 sequences were fused either at the N- or C-termini, based on steric considerations (RBD-SpyT2, SpyT2-NP, Spike-SpyT2). Conjugation of stabilised and trimeric Spike-SpyT2 to the dodecameric SpyC-Dps leads to polymerisation due to the multivalency of both partners. To overcome this problem, and to obtain a biochemically defined sample, we co-transfected HEK 293T Lenti-X cells with two different plasmids in a 3 to 1 ratio, one expressing a SpyT2 version and one without SpyT2. This favoured the expression of Spike trimers in which only one of the monomers contains the SpyTag. Stabilised, trimeric and on average monovalent Spike-SpyT2 and also RBD-SpyT2 were purified from conditioned media of HEK 293S GnT1^-/-^ (for Spike-SpyT2) or Expi 293 (for RBD-SpyT2) cell cultures. SpyC-Dps and SpyT2-NP were purified from the cytosol of *E. coli* cells transformed with the appropriate plasmids. All constructs possess histidine tags and were purified by immobilised metal affinity chromatography (IMAC) and at least one additional size exclusion step (SEC). Sequences of all proteins used can be found in Suppl. Table 1. Expression yields were excellent in all cases: SpyC-Dps yielded ∼120 mg/L culture, RBD-SpyT2 ∼40 mg/L culture, stabilised trimeric and monovalent Spike-SpyT2 ∼13 mg/L culture and SpyT2-NP ∼60 mg/L culture of pure proteins (Fig. 2A).

**Figure 2:**
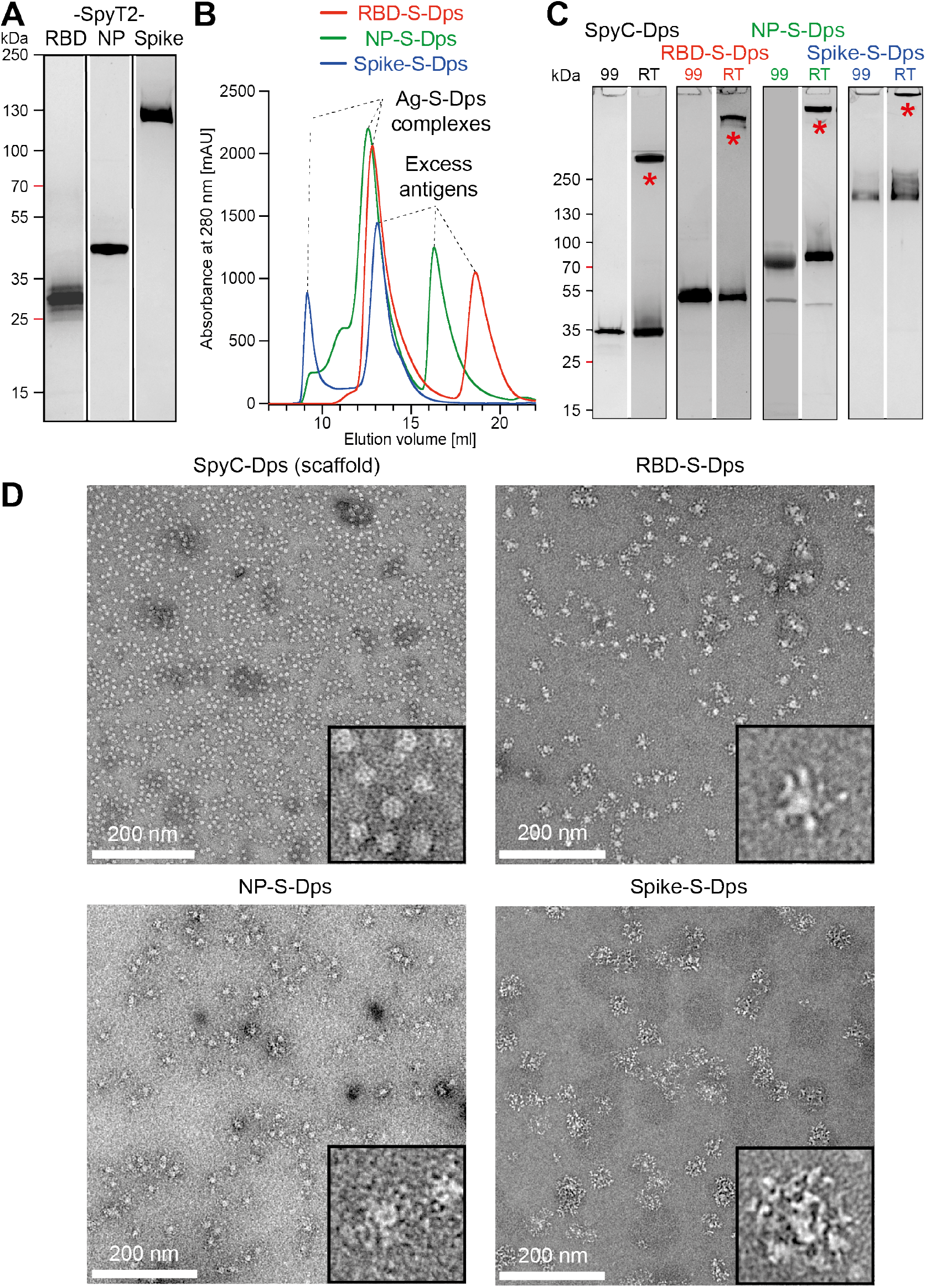
Preparation and quality control of coupled antigen – Dps complexes (Ag-S-Dps). **A**) SDS-PAGE of the three expressed and purified antigens as introduced in Fig. 1C. Glycosylation of Spike leads to a fuzzy appearance of its band. RBD-SpyT2 and Spike-SpyT2 were expressed in mammalian cells, SpyT2-NP was expressed in bacteria, as was the SpyC-Dps scaffold. **B**) Size-exclusion chromatography to separate excess antigens after the SpyCatcher/Spytag2 coupling reactions; Superose 6 Increase in PBS. **C**) SDS-PAGE of the coupled and purified Ag-S-Dps complexes. “RT”, no heating; “99”, heated to 99°C. The SpyC-Dps scaffold alone, as well as all the three coupled complexes show high-molecular weight complexes, presumably dodecameric, that disappear only after heating of the samples in SDS loading buffer. **D**) Negative-stain electron microscopy analyses of the three multimeric Ag-S-Dps complexes, showing that all samples form defined and monodisperse spheres that display the antigens on their surface, leading to particles of different sizes for the three differently-sized antigens.

To achieve efficient coupling of scaffold and antigens, a molar excess of each of the three purified antigens (RBD-SpyT2, SpyT2-NP, Spike-SpyT2) was mixed with SpyC-Dps to facilitate covalent coupling. Subsequent removal of excess antigens was accomplished by SEC using a Superose 6 column (Fig. 2B). Coupling efficiency was analysed by SDS-PAGE, followed by Coomassie staining (Fig. 2C). When the coupled samples were mixed with denaturating SDS sample buffer without additional heating, we detected high molecular weight complexes that we suggest represent dodecameric assemblies caused by Dps that survive SDS treatment (“RT” lanes). Heating the samples to 99 °C led to the disappearing of the higher bands (“99” lanes), confirming both the (SDS-) stability and the purity of the coupled and multimerised protein samples. Note that there were no bands showing uncoupled SpyC-Dps in any of the three Ag-S-Dps samples, meaning that coupling used all 12 available Dps subunits.

Next, we analysed the integrity of the scaffold after the coupling reactions, as well as homogeneity by electron microscopy (Fig. 2D). For the scaffold alone, SpyC-Dps, we observed the expected small and well-dispersed ∼10 nm Dps spheres. Similar homogeneity and monodispersity were observed for all three coupled Ag-S-Dps versions, RBD-S-Dps, NP-S-Dps and Spike-S-Dps. The Ag-S-Dps complexes were larger than the scaffold alone as the Dps spheres were densely surrounded by extra densities, indicating the success of the coupling and the structural integrity of Ag-S-Dps complexes after the coupling reactions. We note that no aggregation was observed for Spike-S-Dps, indicating that the co-transfection approach produced mostly trimeric Spike proteins with only one SpyTag2 present. Taken together, we showed that the scaffold and the three antigens could be produced easily and at high yields and resulted in biochemically pure and defined Ag-S-Dps proteins that display 12 antigens on each Dps scaffold.

To determine whether the coupled Ag-S-Dps complexes were stable in blood plasma for immunisations, we mixed the RBD-S-Dps complex with human serum (clotted, not heat inactivated, at a 1:3 volume ratio). The RBD-S-Dps complex was remarkably stable, with 50% remaining intact after 37 h at 37 °C (Suppl. Fig. 1A & B). Given the stability of the Dps scaffold both in serum and when exposed to denaturing conditions (SDS-PAGE, “RT” lane) (Fig. 2C), we next investigated whether the coupled RBD-S-Dps sample would survive lyophilisation and subsequent re-solubilisation. A lyophilised, dry sample would facilitate prolonged storage even in the absence of refrigeration. We therefore freeze-dried RBD-S-Dps and after rehydration found no evidence of precipitation or significantly reduced protein concentration by SDS-PAGE (Suppl. Fig. 1C). There was also no disappearance of the SDS-stable high-molecular weight band, indicating Dps sphere integrity was maintained after re-hydration. Finally, electron microscopy analysis showed the rehydrated sample to be indistinguishable from the starting material with no evidence of disintegration or aggregation (Suppl. Fig. 1D).

### Multimerisation by Dps greatly enhances immunogenicity, especially for RBD

Having obtained the three multimerised antigen-Dps (Ag-S-Dps), we tested whether they induce a stronger immune response than their monomeric equivalents. We immunised mice with the following protocol: five male C57BL/6J mice per group were given 50 µg protein subcutaneously on days zero and 23, and 25 µg on day 64 (using CpG 1668 as an adjuvant) (Fig. 3A). Blood samples were taken on days 13 (1^st^ bleed), 34 (2^nd^ bleed) and 74 (3^rd^ bleed). After the 1^st^ boost on day 34, antigen-specific antibodies were detected in the sera from the mice by ELISA (Fig. 3B). Substantially higher antibody titres were detected with multimerised Dps-fused RBD and NP. Multimerisation only improved Spike titres modestly, which may be expected given that Spike is already a trimer without Dps. After 74 days, and the second boost, the specific antibody titres were further increased. Spike induced the weakest response and multimerisation had the smallest effect. In contrast, RBD-S-Dps and NP-S-Dps induced substantial increases in antibody titres compared to the non-multimerised versions. We also analysed sera for antibodies against the scaffold protein itself (SpyC-Dps). Sera from mice immunised with coupled Ag-S-Dps complexes showed measurable but low antibody titres against SpyC-Dps. Anti-SpyC-Dps responses remained low even after the second boost, suggesting that the scaffold itself is poorly immunogenic and that in the context of the fusions the antibody response is largely directed against the viral antigens displayed on the surface. Taken together the data show that multimerised Ag-S-Dps complexes produce substantial improvements in antibody titres over the uncoupled antigens. Overall, the strongest response was observed for RBD-S-Dps.

**Figure 3:**
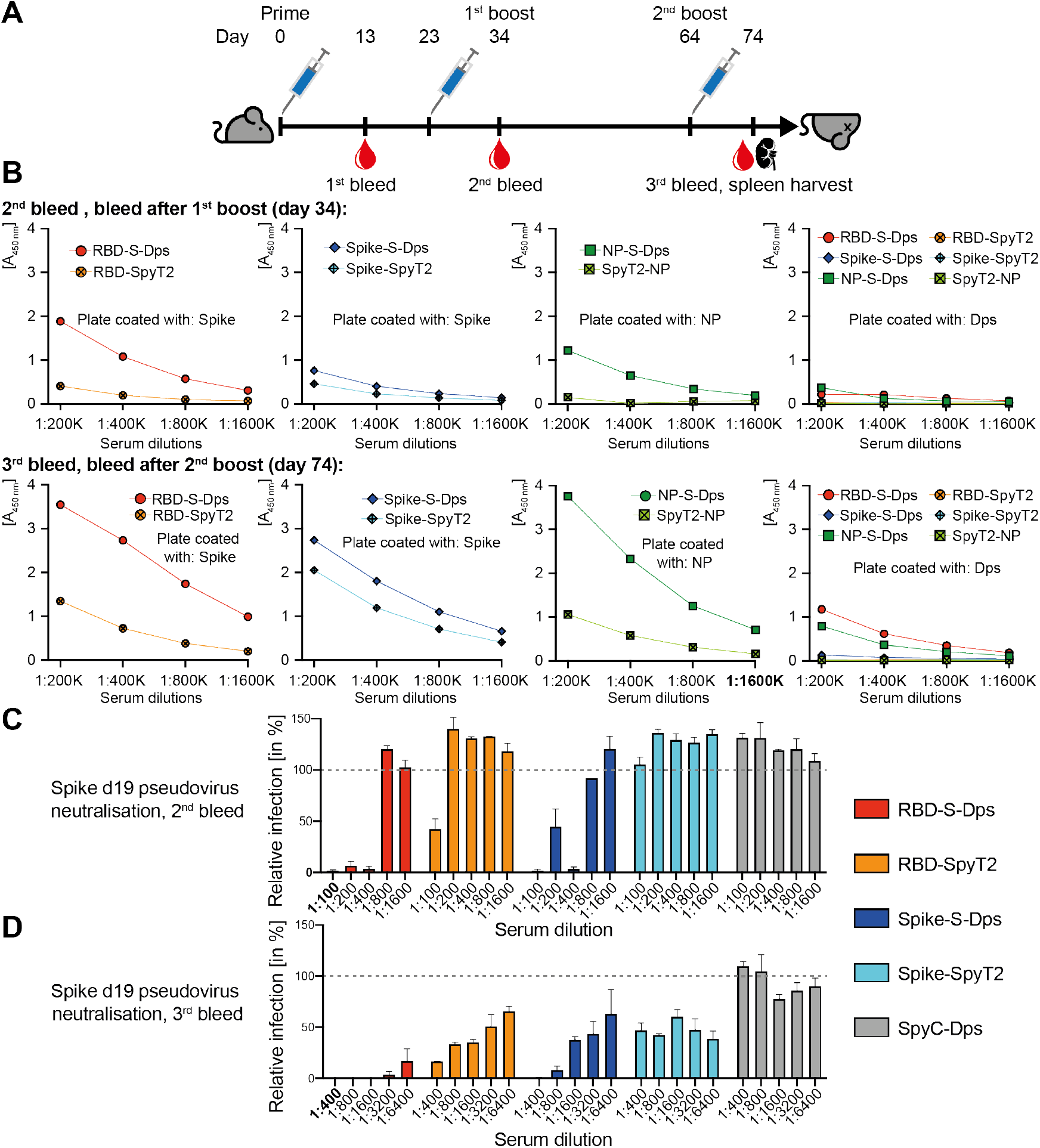
Mouse immunisation – multimeric Ag-S-Dps complexes elicit a powerful and neutralising antibody response in mice. **A**) Immunisation protocol. **B**) Bleeds on day 34 and 74 were tested for binding activity by ELISA against Spike-SpyT2, NP-SpyT2 or polymeric scaffold, SpyC-Dps. In all cases did the multimerised Ag-S-Dps complexes produce more antibodies than the non-multimerised versions. RBD-S-Dps and NP-S-Dps produced very strong responses. **C**) Pseudoviral cell entry neutralisation assay with sera from the 2^nd^ bleed. Sera from immunised mice were tested for neutralisation activity against a Spike-pseudotyped lentiviral GFP vector (hence NP-S-Dps sera will not neutralise). Relative infection is plotted 72 hrs after vector addition by quantifying GFP expression in HEK 293T ACE2/TMPRSS2 target cells. The multimerised RBD-S-Dps and Spike-S-Dps showed strong neutralisation, in contrast to their non-multimerised precursors. **D**) Same as C) but using sera from the 3^rd^ bleed. Neutralisation activity is enhanced in all sera, and the differential between multimerised and non-multimerised antigens remains. Overall, RBD-S-Dps showed the strongest neutralisation activity.

Next, we tested the neutralisation activity of antibodies produced by the mice immunised with RBD-S-Dps, RBD-SpyT2, Spike-S-Dps, and Spike-SpyT2. The mouse sera within each group were pooled at day 34 (2^nd^ bleed) or 74 (3^rd^ bleed) and analysed using a pseudovirus infection assay (note that NP-directed sera will not have an effect in this assay because pseudotyped viruses do not contain NP). In this assay, a lentiviral vector expressing GFP is pseudotyped with Spike protein from SARS-CoV-2 to obtain virions that display Spike in their envelope and infect cells in an ACE2-dependent manner. As seen in Figure 3C, the day 34 sera pool of the multimerised RBD-S-Dps group protected against pseudovirus infection up to a dilution of 1:400, whereas the monomeric RBD-SpyT2 only showed a protective effect at a 1:100 dilution, and even then only reduced infection by ∼50%. Sera from mice immunised with multimeric Spike-S-Dps also protected against infection, whilst Spike-SpyT2 sera were unable to neutralise at any of the dilutions tested. The sera taken after 74 days had substantially increased neutralisation activity (Fig. 3D). The sera from RBD-S-Dps-immunised mice gave the strongest protection: even at a 1:6400 dilution only ∼10% infection could be detected. At this 1:6400 dilution, the monomeric RBD-SpyT2 and Spike-S-Dps sera gave very little neutralisation. While pseudoviruses are widely used to test the neutralisation activity of SARS-CoV-2 antisera, they are based on a lentiviral rather than coronavirus particle and do not recapitulate live virus replication. We therefore tested whether antibodies raised against multimeric RBD-S-Dps were capable of blocking a spreading infection of a primary clinical isolate of SARS-CoV-2. Viral replication was measured by RT-qPCR using probes against *NP* (gRNA) or *E* (sgRNA). RBD-S-Dps antisera from five different animals all potently inhibited SARS-CoV-2 (Suppl. Fig. 2A & B). In contrast, the potency of antisera raised against RBD-SpyT2 varied considerably between mice. We conclude that immunisation with RBD-S-Dps not only produces the highest titre antibodies (Fig. 3B), but also the most neutralising (Fig. 3C & D) and with reliable potency against live virus (Suppl. Fig. 2A & B).

### Single-shot immunisation with multimerised RBD-S-Dps protects mice against SARS-CoV-2 infection

Encouraged by these results, we wanted to know if antigen display on our Dps scaffold would induce a sufficiently strong antibody response to protect animals from SARS-CoV-2 infection. We selected our most potent immunogen, RBD-S-Dps, and used it to immunise mice transgenic for human ACE2 (K18-hACE2) (Zheng et al., 2021). As a single dose vaccination regime offers many downstream logistical and practical benefits, we opted to immunise only once and then challenge with SARS-CoV-2 on day 28 (Fig. 4A). We immunised subcutaneously six K18-hACE2 mice with RBD-S-Dps, six with RBD-SpyT2 and six with PBS (always three female and three male mice), each with 25 µg of the immunogens (except PBS control), plus CpG adjuvant. The anti-Spike antibody response following immunisation was measured by ELISA on days 13 and 24 (before challenge) and on day 35, (seven days post-challenge). A strong anti-Spike antibody titre was detected in RBD-S-Dps-immunised mice, but almost none for either RBD-SpyT2 or PBS (Fig. 4D). Antibody titres remained high for RBD-S-Dps at days 24 and 35.

**Figure 4:**
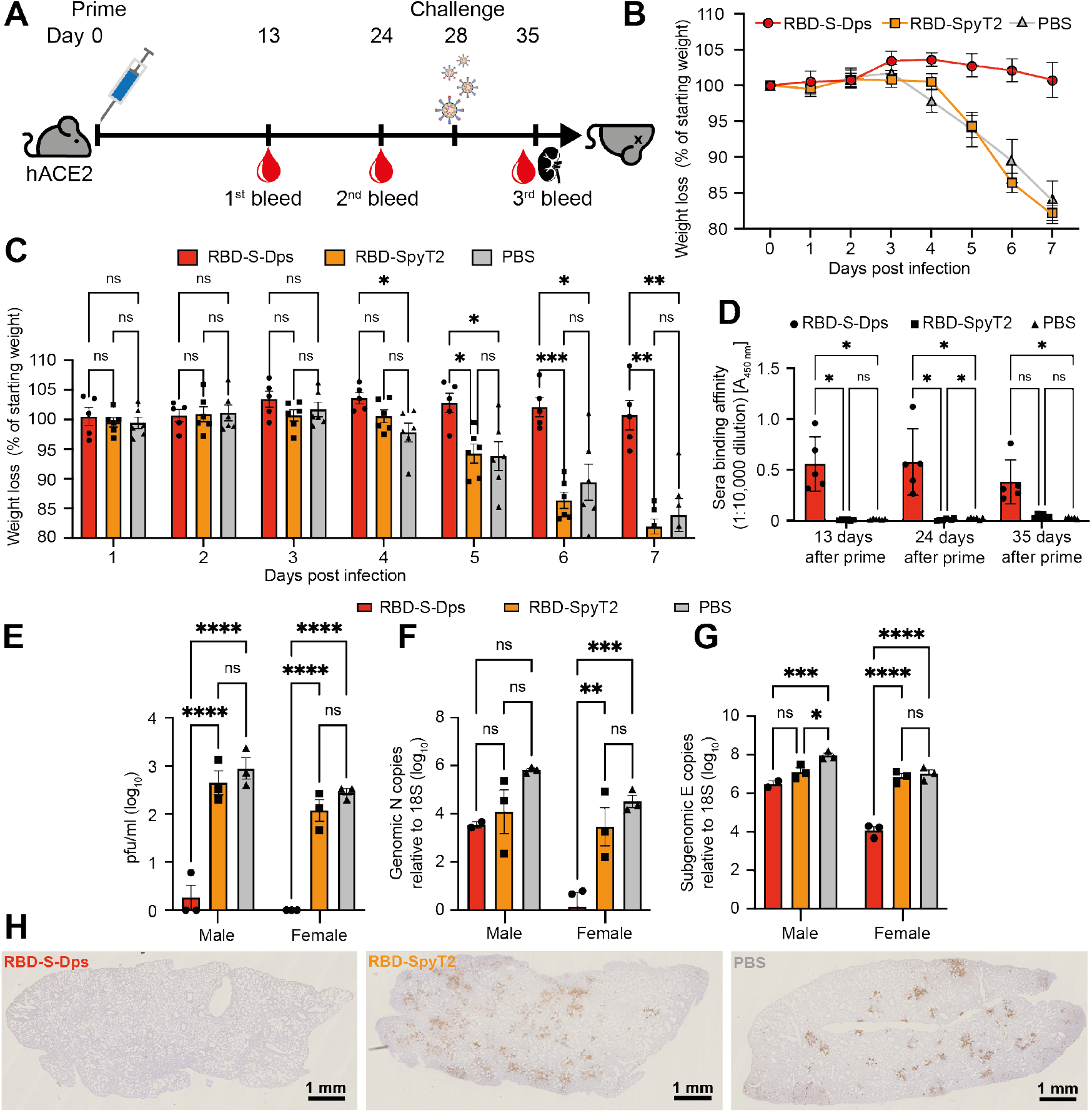
Single-shot immunisation and Sars-CoV-2 challenge experiment using hACE2-mice. **A)** Immunisation and challenge protocol. **B**) K18-hACE2 mice were immunised with 25 µg of RBD-S-Dps, RBD-SpyT2 or given PBS, plus 10 µg CpG adjuvant. The animals were challenged on day 28 with 10^4^ PFU SARS-CoV-2 and changes in weight recorded. The animals in the PBS control group and those who had been given RBD-SpyT2 showed the characteristic weight loss after four days post infection. RBD-S-Dps-immunised mice showed no such weight loss. **C**) Two-way ANOVA tests on the weight changes between groups, as plotted in B). **D**) Sera from days 13, 24 and 35 were tested for anti-RBD antibodies by ELISA. Only RBD-S-Dps mice showed significant antibody. **E**) Plaque assay using lung homogenates from mice culled seven days post-infection. RBD-S-Dps-immunised mice contained very low amounts of infectious SARS-CoV-2 in their lungs. **F) and G**) Genomic and subgenomic (gRNA, sgRNA) qPCR on RNA extracted from lung homogenates, using probes against NP or E, respectively. Two-way ANOVA tests were carried out with significance levels of: p = < 0.05 (*), p = < 0.05 (**), p = < 0.005 (***), p = < 0.0005 (****). **H)** Representative lung sections from animals (n=6) taken seven days post-challenge, stained by immunohistology for SARS-CoV-2 NP protein.

On day 28, animals were challenged with 10^4^ PFU SARS-CoV-2. Mice in the PBS control and RBD-SpyT2-immunised groups began to show clinical signs of illness and a decline in body weight from day four post-infection (Fig. 4B), consistent with previous reports of infection in naïve animals (Zheng et al., 2021). In contrast, mice immunised with multimerised RBD-S-Dps maintained body weight until the day seven end point. There was a statistically significant difference in weights between the RBD-S-Dps-immunised and PBS control groups from day four, and between RBD-S-Dps- and RBD-SpyT2-immunised mice from day five (Fig. 4C). There was no significant difference in weight loss between the RBD-SpyT2-immunised mice and PBS controls at any time point, suggesting that, unlike RBD-S-Dps, non-multimerised RBD does not provide protection after only a single vaccination. All animals were culled on day seven post-infection and tissues collected for analysis. As mentioned, there were no significant changes in anti-Spike antibody levels pre-versus post-challenge, indicating that mostly antibodies raised during the immunisation contributed to the immune response during the infection (Fig. 4D). SARS-CoV-2 infection of the lung was quantified by plaque assay and genomic and subgenomic qPCR, using probes against the viral genes NP and E, respectively. There were significantly lower levels of infectious virus in the lungs of mice immunised with RBD-S-Dps, compared to either RBD-SpyT2 immunised or PBS control groups (Fig. 4E). A broadly similar pattern was observed when quantifying virus by either genomic or subgenomic qPCR (Fig. 4F & G). However, we noted a marked difference in the amounts detected between male *vs* female mice. Female RBD-S-Dps-immunised mice had significantly fewer genomic and subgenomic transcripts, compared to mice from other groups and their male equivalents (Fig. 4F & G and Suppl. Fig. 3A & B). We attempted to correlate this with differences in antibody titres, but while there was a trend towards lower titres in male mice, particularly just before and just after the challenge, this was not significant (Suppl. Fig. 3C). A larger group size would be needed to confirm this result. Despite these sex-dependent differences in the qPCR data, the near-absence of infectious virus in both male and female RBD-S-Dps immunised mice, as measured by plaque assay (Suppl. Fig. 3D), suggests they were both highly protected. Finally, we examined the lungs of mice from the different groups for histopathological changes and for viral antigen expression using an anti-NP antibody to reveal sites of replication (Fig. 4H, Suppl. Fig. 4) and immune cell infiltration (Suppl. Fig. 5). A detailed description is provided in the Supplementary Results. In summary, lungs from RBD-SpyT2-immunised mice or PBS control mice showed substantial and wide-spread NP expression mainly in pneumocytes (Fig. 4H & Suppl. Fig. 4), indicative of viral replication throughout the lobe and consistent with the high virus levels measured in these animals (Fig. 4E-G). There was also evidence of pneumocyte degeneration and syncytial cell formation, as has been reported in COVID-19 cases post-mortem (Bussani et al., 2020). Multifocal leukocyte infiltration was observed, particularly in PBS-control animals, dominated by macrophages, followed by T-cells (mainly CD4+ and lesser CD8+ cells), B cells and neutrophils (Suppl. Fig. 4 & 5). This is reminiscent of the hyperinflammation in post-mortem reports of lethal COVID-19 associated with immunopathology (Schurink et al., 2020). In contrast, the lungs of mice protected by multimerised RBD-S-Dps were either almost or entirely clear of NP expression (Suppl. Fig. 4) and pathological changes (female mice), or showed only mild changes consistent with those observed in the PBS-control animals (Suppl. Fig. 5), and markedly reduced NP expression. Taken together, these data indicate that immunisation with RBD-S-Dps is highly protective against SARS-CoV-2 in hACE2-expressing mice, even after a single dose, whilst monomeric RBD-SpyT2 is not.

## DISCUSSION

Here we have shown that the ferritin-like protein Dps from the hyperthermophile *S. islandicus* possesses exceptional qualities as a SARS-CoV-2 subunit vaccine scaffold. We combined Dps with the SpyCatcher/SpyTag system in order to create a “plug-and-play” system that allows the rapid and facile synthesis of highly stable multimeric subunit vaccines. Mixing the SpyCatcher-Dps protein with any compatible SpyTag antigen leads to the assembly of highly monodisperse nanoparticles displaying exactly 12 antigens. Using this approach, we have produced subunit vaccines based on Spike, RBD or NP from SARS-CoV-2 and tested them in immunisation and viral challenge experiments. In each case, the Dps-displayed antigens out-performed their non-multimerised equivalents and induced a more rapid and potent antibody response.

Subunit vaccines offer distinct advantages in cost, simplicity, production capacity, storage, transport and administration over nucleic-acid based vaccines (Pollet et al., 2021). Principle amongst these considerations is stability, with currently used vaccines such as those produced by Pfizer–BioNTech, Moderna and Oxford-AstraZeneca requiring a -80°C or -20°C cold-chain. In countries without a highly developed logistical and medical infrastructure this represents a significant impediment to vaccination. Whilst subunit vaccine development currently lags behind nucleic-acid based equivalents, there is good evidence that such vaccines are nevertheless effective at inducing a protective response. SARS-CoV-2 RBD by itself (Yang et al., 2020) or in simple fusions such as to IgG Fc (Liu et al., 2020) have been shown to elicit SARS-CoV-2-neutralising antibodies. Antigen multimerisation increases neutralising titres, for instance when using ferritin as a scaffold (He et al., 2021; Powell et al., 2021). More complex scaffolds have also been used, for instance virus-like icosahedral particles that display 60 antigen copies (e.g. I3-01) (Hsia et al., 2016). When fused directly to viral antigens (Walls et al., 2020), or using the SpyCatcher/SpyTag system (Cohen et al., 2021; Tan et al., 2021) the I3-01 scaffold has been shown to induce a neutralising antibody response. Our scaffold differs from those previously used to deliver SARS-CoV-2 immunogens in several important aspects. First, because we have used a thermostable protein it is intrinsically more stable, providing potential benefits to vaccine transport and storage and also to immunogen stability *in vivo*. Second, it is smaller than other scaffolds (< 10 nm vs > 10 nm for ferritin or 25 nm for the I3-01 nanoparticle), making it an easier cargo for cellular uptake. Third, it displays fewer copies than ferritin or I3-01 (12 vs 24 or 60, respectively), allowing the selection of higher-affinity B cells and avoiding the activation of off-target (and possibly cross-reactive) B cell competitors (Kato et al., 2020). Fourth, in contrast to *bona fide* ferritin scaffolds, the N- and the C-termini of Dps are both accessible on the outside of the sphere. This allows, at east in principle, for the conjugation of two discrete antigens onto a single scaffold, for example both SARS-CoV-2 Spike/RBD and NP to be displayed simultaneously.

Importantly, we have provided here data demonstrating the benefit of antigen multimerisation in inducing not just neutralising antibodies but an immune response capable of providing *in vivo* protection. In our SARS-CoV-2 challenge experiments, we found that RBD alone failed to protect mice, which displayed continued weight-loss and high viral loads in the lungs. In contrast, our Dps-based vaccine displaying RBD completely protected mice from SARS-CoV-2-associated pneumonia and disease after only a single immunisation. We noted however a difference in Dps-RBD induced protection between male and female mice, with the latter having lower viral loads and hardly any pulmonary changes. Trial data for both mRNA and vector-based vaccines has not been disaggregated by sex but data on SARS-CoV-2 infection show that men are more at risk of severe adverse conditions, hospitalisation and death (Klein et al., 2020; Scully et al., 2020). Our results support the consideration of sex as a variable in vaccine trials (Bischof et al., 2020).

Further research is needed to develop the Dps-scaffold into a *bona fide* vaccine for SARS-CoV-2 and other viruses. Replicating the robust neutralising antibody response and high-level of protection achieved in mice from a single dose in humans will be crucial. Moreover, whilst most studies of subunit vaccines have focused on antibodies, long-lasting protection is likely to be dependent upon stimulating CD4+ and CD8+ T cell immunity (Sauer and Harris, 2020; Zuo et al., 2021). Data from current vaccine trials and roll-outs has yet to be fully analysed but a correctly balanced T cell response appears associated with recovery from acute infection and the avoidance of hospitalisation and severe virus-induced immunopathology (Chen and John Wherry, 2020). Fortunately, the analysis of T cell epitopes from SARS-CoV-2 convalescents (Nelde et al., 2021) provides a basis for engineering subunit vaccines specifically to engage both B and T cells. In this context, the ability of our Dps-scaffold to display antigens at both termini may prove particularly beneficial. In addition to ensuring a well-balanced immune response in humans, a more thorough investigation into the long-term stability, storage and reconstitution of lyophilised material is required to demonstrate that a Dps-based vaccine is suitable for use in regions with limited infrastructure. Future work notwithstanding, our data add to a body of evidence that subunit-based vaccines represent a viable choice as a vaccine modality for SARS-CoV-2. Whilst other vaccine formats are significantly more advanced, subunit approaches such as Dps offer distinct advantages in simplicity of production, requiring no proprietary technology, robustness of material and potency of protection.

## MATERIAL & METHODS

### Cloning, expression and purification of the protein components

#### SpyC-Dps

a hexa-histidine tag was fused to ΔN1-SpyCatcher, which was subsequently linked to the Dps from *Sulfolobus islandicus* (ORF SIL_0492, GenBank AGJ61963.1), separated by a GSEGSSGG-linker (Suppl. Table 1, SpyC-Dps). The sequence was codon optimised for the expression in *E. coli* and the gene was cloned into the pOPINS vector by Gibson assembly. The plasmid encoding for SpyC-Dps was transformed into C43(DE3) *E. coli*. Cells were grown at 37°C in 2xYT medium to an OD_600_ of 0.8. Protein production was induced with 1 mM IPTG for 6 h. Cells were harvested at 4500 x g for 25 min at room temperature (RT). Cells were shock-frozen in liquid nitrogen (LN2) and stored at -80 °C. Cells producing SpyC-Dps were re-suspended in T-buffer1 (30 mM Tris, 250 mM NaCl, pH 8.0) with one tablet of Complete Protease Inhibitors (Roche) per 10 g cells wet weight. Cell disruption was carried out using sonication for 7.5 min “on” time, using a 50 % duty cycle. Cell debris were removed by centrifugation at 20,000 x g for 30 min at RT. The supernatant was loaded onto a HisTrap FF affinity chromatography column (Cytiva). Washing was carried out for 17 column volumes (CV) with T-buffer1 plus 110 mM imidazole. The protein was eluted with T-buffer1 containing 400 mM imidazole. Purity of fractions was examined by SDS-PAGE and the purest fraction were pooled and concentrated using a Vivaspin Turbo centrifugal concentrator (100,000 MWCO, Sartorius). Concentrated sample was loaded onto a size-exclusion column (SEC, Sephacryl S-400, Cytiva), with PBS as the running buffer. Purity was examined by SDS-PAGE and the sample was frozen in LN2 and stored at -80 °C.

#### SpyT2-NP

the nucleocapsid protein (amino acids 48 - 364; GenBank: MN908947; NP) was cloned into the vector pOP-TH and N-terminally equipped with a hexa-histidine tag (Pickering et al., 2020). A SpyTag2 sequence separated by GS-rich linkers was inserted between the hexa-histidine tag and NP (Fig. 1 & Suppl. Table 1, SpyT2-NP). The vector encoding for SpyT2-NP was transformed into *E. coli* C41(DE3) cells. For protein expression, cells were grown at 37 °C in 2xYT medium to an OD_600_ of 0.7. Protein production was induced with 1 mM IPTG for 6 h. Cells were harvested at 4500 x g for 25 min at 4 °C. Cells were frozen in LN2 and stored at -80 °C. SpyT2-NP-producing cells were re-suspended in T-buffer2 (50 mM Tris, 1 M NaCl, 10 mM imidazole, 2 mM DTT, pH 8.0) with Complete Protease Inhibitor added (1 tablet per 10 g cells wet weight). Cells were lysed by sonication (3 mins total “on” time, duty cycle 50 %). Precipitated proteins and cell debris were removed by centrifugation (40,000 x g, 1 h, 4 °C). The supernatant was loaded onto a HisTrap FF affinity chromatography column and washed with 20 CV T-Buffer3 (50 mM Tris, 300 mM NaCl, 1 mM DTT, pH 8.0) containing 20 mM imidazole. Elution was carried out in T-buffer3 containing 400 mM imidazole. Elution fractions containing NP were loaded onto 20 ml HiTrap Heparin HP column equilibrated in T-buffer4 (50 Tris, 1 mM DTT, pH 8.0). The column was washed with 3 CV T-buffer4. Elution was carried out with a linear gradient of 0 - 2 M NaCl. Elution fractions containing SpyT2-NP were examined by SDS-PAGE and pooled, and concentrated using a Vivaspin Turbo concentrator with a 10,000 MWCO (Sartorius). Concentrated sample was loaded onto a SEC column (Sephacryl S-200) (Cytiva) in PBS + 250 mM additional NaCl. Purity was checked by SDS-PAGE and samples were frozen in LN2 and stored at -80 °C.

#### Spike-SpyT2 and Spike

to express the ectodomain of the stabilised prefusion Spike protein trimer (Wrapp et al., 2020) with only one subunit carrying the SpyTag2 tag, two constructs – one with and one without a SpyTag2 - were made. First, a gene encoding residues 16-1208 of SARS-CoV-2 Spike protein (GenBank: MN908947) with proline substitutions at residues 986 and 987, a GSAS substitution at the furin cleavage site (residue 682-685), a C-terminal T4 fibritin trimerisation motif, a GGSGGS linker, an HRV3C protease cleavage site, a GGS linker and an AviTag, was synthesised and cloned into the lentiviral expression vector pHR-SFFV (Chang et al., 2015, 2016; Elegheert et al., 2018) downstream of the sequence encoding the chicken RPTPσ secretion signal peptide (cRPTPσSP) (Aricescu et al., 2006). Then, either a GGS linker and a hexa-histidine tag, or a GGS linker, an octa-histidine tag, a GGSGGSGGS linker and a SpyTag2 were inserted after the AviTag sequence to form two Spike constructs, with and without a SpyTag2 (Suppl. Table 1, Spike-SpyT2 and Spike, respectively). For protein expression and purification, please see the next paragraph.

#### RBD-SpyT2

a gene encoding residue 332-529 of SARS-CoV-2 Spike protein (constituting the receptor binding domain, RBD) was synthesised and cloned downstream of cRPTPσ of the pHR-SFFV vector and a GGSGGS linker, an AviTag, a GGS linker, an octa-histidine tag, a GGSGGSGGS linker and a SpyTag2 were inserted at the 3’ end of the gene (Suppl. Table 1, RBD-SpyT2). The vectors for Spike-SpyT2, Spike and RBD-SpyT2 were used for protein production in the mammalian lentiviral expression system (Chang et al., 2015, 2016; Elegheert et al., 2018). The DNA of the constructs was mixed with the lentiviral envelope and packaging vectors pMD2-G and psPAX2c (Addgene) and polyethylenimine (PEI, Sigma) to transiently transfect HEK 293T Lenti-X cells (Takara/Clontech) to make lentiviral particles. To make Spike trimer protein with only one subunit carrying a SpyTag2, the DNAs of constructs Spike and Spike-SpyT2 were used at a molar ratio 3:1. The virus particles produced were used to infect HEK 293S GnT1-/- cells (for Spike-SpyT2) or Expi 293 cells (for RBD-SpyT2). The infected cells were then expanded to obtain 3 L cultures and conditioned media were harvested and sterile filtered (0.22 μm). The supernatant was concentrated and the buffer exchanged to 25 mM Tris pH 8.0, 300 mM NaCl using an Äkta flux tangential flow system (Cytiva). The conditioned supernatant was then loaded onto a HisTrap column (Cytiva) and washed and eluted with 50 mM and 250 mM imidazole in the same buffer, respectively. Eluted fractions were checked by SDS-PAGE, pooled and further purified in PBS buffer by SEC on Superdex 200 for RBD and Superose 6 for trimeric Spike protein (both Cytiva). Peak fractions were checked by SDS-PAGE again and frozen in LN2 and stored at -80 °C.

### Coupling and purification of multimerised complexes

For the preparations of Ag-S-Dps complexes, comprising RBD-S-Dps, NP-S-Dps and Spike-S-Dps the antigens: RBD-SpyT2, SpyT2-NP and Spike/Spike-SpyT2, and the scaffold protein SpyC-Dps were diluted in PBS buffer + 250 mM NaCl to 0.2 to 1 mg/mL and mixed. To achieve full occupancy of SpyC-Dps with the antigens, the molar ratio for SpyC-Dps to RBD-SpyT2 was 1:1.3, for SpyT2-NP 1:2 and for trimeric Spike/Spike-SpyT2 1:2.5. Reactions were left for ∼5 min at RT and covalent coupling between SpyCather2 and SpyTag2 was checked by SDS-PAGE. Subsequently, samples were concentrated using Vivaspin Turbo concentrators (100,000 MWCO). Antigen-decorated SpyC-Dps complexes were separated from the excess antigens by SEC in PBS + 250 mM NaCl on a Superose 6 Increase column (Cytiva). Fractions were checked again for purity by SDS-PAGE, frozen in LN2 and stored at -80 °C.

### Negative-stain electron microscopy

Proteins were diluted in PBS to concentrations of ∼0.012 mg/mL. 3 µL of the solution were applied to a glow-discharged carbon-coated grid and immediately blotted. For the staining, 10 μL of 2% (w/v) uranyl formate were applied and removed immediately by blotting the grid with filter paper. Images were collected on a FEI Tecnai Spirit 120 kV electron microscope, equipped with a CCD detector.

### *In vitro* human plasma stability assay

The *in vitro* stability of RBD-S-Dps was studied in clotted human plasma (MD Biomedicals, cat. #2930149). Stocks of the RBD-S-Dps samples (751.7 kDa per dodecameric complex) were diluted in PBS to a final concentration of ∼0.8 µM and subsequently mixed with pre-warmed human plasma in a 1:3 (protein:plasma, v/v) ratio. The mixtures were incubated at 37 °C for seven days. Samples were taken after 0, 1, 24, 48, 72, 96, 120 and 168 h, and immediately mixed with denaturing gel-loading buffer, followed by 30 min incubation at 99 °C. Inactivated samples were stored at -20 °C before the samples were diluted 1:10 with 1x SDS sample buffer and 5 µL per sample were analysed by SDS-PAGE and Western blotting. The Ag-S-Dps complexes were detected using the HisProbe-HRP (Thermo Fisher Scientific, TFS) and human transferrin was used as a loading control and detected using transferrin antibodies from chicken and chicken-HRP conjugated antibodies (Thermo Fisher Scientific, TFS, cat. #PA1-9525 and cat. # 31401).

### Lyophilisation of samples

An aliquot of RBD-S-Dps of 120 µL (at a protein concentration of 1.4 mg/mL, in PBS buffer plus additional 250 mM NaCl) was divided into a 40 µL control and a second aliquot of 80 µL. The 80 µL aliquot was lyophilised for 4 h at 30 °C with the aid of a vacuum concentrator (Eppendorf Concentrator 5301) attached to a refrigerated condensation trap (Savant). After lyophilisation to complete dryness, the sample was resuspended in 80 µL Milli-Q water. The sample was not centrifuged or processed in any other way after rehydration. EM grids were prepared by staining 1:20 and 1:100 dilutions (in PBS plus 250 mM NaCl) of lyophilised and resuspended sample with 2 % uranyl formate solution on carbon-coated CF400-CU-UL grids (Electron Microscopy Sciences) as described earlier. Imaging was also performed as described earlier. 10 µL of lyophilised and rehydrated sample and the untreated control were compared by SDS-PAGE followed by Coomassie staining.

### Mouse immunisation (Fig. 3)

Six weeks-old C57BL/6J mice (Jackson) were used in immunisation experiments, which were conducted in accordance with the E7 moderate severity limit protocol and the UK Home Office Animals (Scientific Procedures) Act (ASPA, 1986), and approved by the UKRI Animal Welfare and Ethical Review Body. Mice were initially (prime) immunised subcutaneously with 50 µg of the antigens in PBS, mixed with 10 µg CpG ODN 1668 adjuvant (InvivoGen). The following antigens were used: RBD-S-DPS, RBD-SpyT2; NP-S-DPS, SpyT2-NP; Spike-S-DPS, Spike/Spike-SpyT2 and SpyC-Dps. Mice were subcutaneously boosted with 50 µg antigens at day 23 and with 25 µg antigens at day 64. Tail bleeds for ELISA analyses were collected on days 13 and 34.

### Preparation of SARS-CoV-2 Spike-pseudotyped HIV-1 virions

Replication deficient SARS-CoV-2 pseudotyped HIV-1 virions were prepared as described previously (Morecroft and Thomas, 1988). Briefly, virions were produced in HEK 293T cells by transfection with 1 µg of the plasmid encoding SARS CoV-2 Spike protein (pCAGGS-SpikeΔc19), 1 µg pCRV GagPol and 1.5 μg GFP-encoding plasmid (CSGW). Viral supernatants were filtered through a 0.45 μm syringe filter at 48 h and 72 h post-transfection and pelleted for 2 h at 28,000 x g. Pelleted virions were drained and then resuspended in DMEM (Gibco).

### Spike-pseudotyped neutralisation assays with mouse sera

HEK 293T-hACE2-TMPRSS2 cells were described previously (Papa et al., 2021). Cells were plated into 96-well plates at a density of 0.75 x 10^3^ cells per well and allowed to attach overnight. 20 µL pseudovirus-containing supernatant was mixed with 2 µL dilutions of heat-inactivated mouse sera and incubated for 40 min at RT. 10 µL of this mixture was added to cells. 72 h later, cell entry was detected through the expression of GFP by visualisation on an Incucyte S3 live cell imaging system (Sartorius). The percent of cell entry was quantified as GFP positive areas of cells over the total area covered by cells. Entry inhibition by the sera was calculated as percent virus infection relative to virus only control.

### ELISA assays

96-well plates (Nunc) were coated overnight with 5 µg/mL of the indicated antigens. Plates were blocked with MPBST: 2 % (v/v) milk in PBS, 0.05 % Tween 20. Polyclonal sera from individual mice (challenge experiment) or mouse sera pooled within the same group (mouse immunisation) were diluted as indicated with MPBST and incubated for 45 min on antigen-coated plates. Plates were washed with MPBST and bound antibodies were detected with goat anti-mouse IgG-HRP (Jackson Immunoresearch, #115-035-071).

### Cell culture and virus

UK strain of SARS-CoV-2 (hCoV-2/human/Liverpool/REMRQ0001/2020; PANGO lineage B), was used and grown to P4 in Vero E6 cells (Patterson et al., 2020). The intracellular viral genome sequence and the titre of virus in the stock was determined by direct RNA sequencing (Genbank: MW041156). The virus stock did not contain a deletion of the furin cleavage that has been described previously during passage (Davidson et al., 2020).

### Mouse SARS-CoV-2 challenge experiment

Animal work was approved by the local University of Liverpool Animal Welfare and Ethical Review Body and performed under UK Home Office Project Licence PP4715265. Mice carrying the human ACE2 gene under the control of the keratin 18 promoter (K18-hACE2; formally B6.Cg-Tg(K18-ACE2)2Prlmn/J) were purchased from Jackson Laboratories. Mice were maintained under SPF barrier conditions in individually ventilated cages. Animals were randomly assigned into multiple cohorts and given 25 µg antigen (RBD-S-DPS or RBD-SpyT2) & 10 µg CpG or PBS via subcutaneous injection. On day 28 post-immunisation, mice were anaesthetised lightly with isoflurane and inoculated intranasally with 50 µL containing 10^4^ PFU SARS-CoV-2 in PBS. They were culled on day 35 post-immunisation by an overdose of pentabarbitone. Tissues were removed immediately for downstream processing.

### RNA extraction and DNase treatment

The upper lobes of the right lung were dissected and homogenised in 1 mL of TRIzol reagent (TFS) using a Bead Ruptor 24 (Omni International) at 2 meters per second for 30 s. The homogenates were clarified by centrifugation at 12,000 x g for 5 min before full RNA extraction was carried out according to manufacturer’s instructions. RNA was quantified and quality assessed using a Nanodrop (TFS) before a total of 1 μg was DNase treated using the TURBO DNA-free kit (TFS) as per manufacturer’s instructions.

### qRT-PCR for viral load

Viral loads were quantified using the GoTaq® Probe 1-Step RT-qPCR System (Promega). For quantification of SARS-COV-2 the nCOV_N1 primer/probe mix from the SARS-CoV-2 (2019-nCoV) CDC qPCR Probe Assay (IDT) were utilised while the standard curve was generated via 10-fold serial dilution of the 2019-nCoV_N_Positive Control (IDT) from 10^6^ to 0.1 copies/reaction. The E sgRNA primers and probe have been previously described (Wölfel et al., 2020) and were utilised at 400 nM and 200 nM respectively. Murine 18S primers and probe sequences were utilised at 400 nM and 200 nM respectively. The IAV primers and probe sequences were published as part of the CDC IAV detection kit (20403211). The IAV reverse genetics plasmid encoding the NS segment was diluted 10-fold from 10^6^ to 0.1 copies/reaction to serve as a standard curve. The thermal cycling conditions for all qRT-PCR reactions were as follows: 1 cycle of 45 °C for 15 min and 1 cycle of 95 °C followed by 40 cycles of 95 °C for 15 s and 60 °C for 1 min The 18S standard was generated by the amplification of a fragment of the murine 18S cDNA using the primers F: ACCTGGTTGATCCTGCCAGGTAGC and R: GCATGCCAGAGTCTCGTTCG. Similarly, the E sgRNA standard was generated by PCR using the qPCR primers. cDNA was generated using the SuperScript IV reverse transcriptase kit (TFS) and PCR carried out using Q5 High-Fidelity 2X Master Mix (New England Biolabs) as per manufacturer’s instructions. Both PCR products were purified using the QIAquick PCR Purification Kit (Qiagen) and serially diluted 10-fold from 10^10^ to 10^4^ copies/reaction to form the standard curve.

### Histology and immunohistology

The left lung lobes were fixed in formal saline for 24 h and routinely paraffin wax embedded. Consecutive sections (3-5 µm) were either stained with haematoxylin and eosin (HE) or used for immunohistology (IH). IH was performed to detect SARS-CoV-2 antigen and leukocyte subtypes, i.e. T cells (CD3+, CD4+, CD8+), B cells (CD45R/B220+) and macrophages (Iba1+), using the horseradish peroxidase (HRP) method and the following primary antibodies: rabbit anti-SARS-CoV NP (Rockland, 200-402-A50), rabbit anti-mouse CD3 (clone SP7; Spring Bioscience Corp.), rabbit anti-mouse CD4 (clone #1; SinoBiological), rabbit anti-mouse CD8 (D4W2Z; Cell Signaling Technology), rat anti-mouse CD45R (clone B220, BD Pharmingen), rabbit anti-human Iba1/AIF1 (Wako, 019-19741). Briefly, after deparaffination, sections underwent antigen retrieval in citrate buffer (pH 6.0; Agilent) (anti-SARS-CoV-2, -CD8, -CD45R, -Iba1) or Tris-EDTA buffer, pH 9.0 (anti-CD3, -CD4) for 20 min at 98 °C and for 20 min at 37 °C respectively, followed by incubation with the primary antibody overnight at 4 °C (anti-SARS-CoV-2), 60 min at RT (anti-CD3, -CD8, -CD45R, -Iba1) or 60 min at 37 °C (anti-CD3, -CD4). This was followed by blocking of endogenous peroxidase (peroxidase block, Agilent) for 10 min at RT and incubation with the secondary antibody, EnVision+/HRP, Rabbit and Rat respectively (Agilent) for 30 min at RT (anti-SARS-CoV, -CD8, -CD45R, -Iba1) or the Omni-Map anti Rb HRP (Ventana) for 16 min at 37 °C (anti-CD3, -CD4), followed by EnVision FLEX DAB+ Chromogen in Substrate buffer (Agilent; anti-SARS-CoV-2, -CD8, -CD45R, -Iba1) for 10 min at RT or the DAB-Map-Kit (Ventana; anti-CD3, -CD4), all in an autostainer (Dako or Ventana). Sections were subsequently counterstained with haematoxylin.

## ACKNOWELDGEMENTS

We thank Mark Wing and Kevin O’Connell (MRC-LMB, Cambridge). We would especially like to thank the biomedical research staff at Liverpool and the LMB’s Ares facility for their help and support, as well as the technical staff of the Histology Laboratory, Institute of Veterinary Pathology, Vetsuisse Faculty Zurich. Work at the MRC-LMB was funded by the Medical Research Council, UK (U105184326 to JL, U105181010 to LCJ, MC_UP_1201/15 and MR/L009609/1 to ARA) and Wellcome, UK (203276/C/16/Z to JL, 200594/Z/16/Z and 214344/A/18/Z to LCJ). Work in Liverpool was funded in by the Medical Research Council, UK (MR/W005611/1 to JPS and JAH) and the US Food and Drug Administration, USA (75F40120C00085 to JAH).

## SUPPLEMENTAL INFORMATION

### SUPPLEMENTAL RESULTS

Detailed description of the lung histology in RBS-D-Dps-immunised mice subsequently challenged with SARS-Cov-2 (Fig. 4A):

#### Mice immunised with PBS control

All animals showed a mild to moderate increase in interstitial cellularity and multifocal extensive areas of consolidation due to macrophage and lymphocyte infiltration, with a few neutrophils and with abundant activated type II cells, occasional syncytial cells and some degenerate cells and moderate mesothelial cell activation above the affected parenchymal area (Suppl. Fig. 4A). The changes were associated with extensive viral antigen expression in type I and type II pneumocytes both in consolidated areas and in alveoli without inflammatory changes. Occasional macrophages in the infiltrate also appeared to express viral antigen (Suppl. Fig. 4B). Macrophages (Iba1+) were the dominant cells in the infiltrates (Suppl. Fig. 5A), followed by numerous T cells with a relatively high proportion and CD4 positive cells and less CD8 positive cells (Suppl. Fig. 5C, E, G), and a moderate number of B cells (Suppl. Fig. 5I). There were also areas where alveoli exhibited type II cell activation and desquamation, with desquamation of alveolar macrophages and some neutrophils in the lumen. In addition, mild to moderate mononuclear vasculitis (mainly arteritis) was seen (Suppl. Fig. 4A). Also, the infiltrate was dominated by macrophages, followed by T cells (CD4 positive cells and less CD8 positive cells) and fewer B cells.

#### Mice immunised with monomeric RBD-SpyT2

All animals showed a mild to moderate increase in interstitial cellularity and multifocal extensive areas of consolidation similar in composition and extent to those seen in the PBS-control mice (Suppl. Fig. 4C). Viral antigen was detected in multiple variably-sized foci, in type I and II pneumocytes and in macrophages within and close to consolidated areas (Suppl. Fig. 4D).

#### Mice immunised with multimerised RBD-S-Dps

The three female animals showed minimal histological changes in the lungs (Suppl. Fig. 4E). Besides a very mild increase in interstitial cellularity, with rare T cells (both CD4 and CD8 positive cells) and B cells, one animal showed scattered small focal leukocyte aggregates. Viral antigen expression was restricted to a few macrophages in the leukocyte aggregates in the latter animal, while a second had positive pneumocyte in an alveolus; in the third lung, viral antigen was not detected (Suppl. Fig. 4F). In male animals, multifocal inflammatory infiltrates with viral antigen expression and mild vasculitis similar to the other two groups, but substantially less extensive were seen (Suppl. Fig. 4G, H). Also here, macrophages were the dominant cells in the focal infiltrates (Suppl. Fig. 5B), followed by the T cells (Suppl. Fig. 5D). The T cell population showed a mild shift in composition, as now CD8-positive cells were as numerous or more frequent than CD4-positive cells (Suppl. Fig. 5F-H). CD8-positive cells were also seen within alveolar lumina (Suppl. Fig. 5B inset). B cells were found in moderate numbers in the infiltrates, in one animal they also formed peribronchiolar aggregates (Suppl. Fig. 5J).

---

**Supplemental Figure 1.**
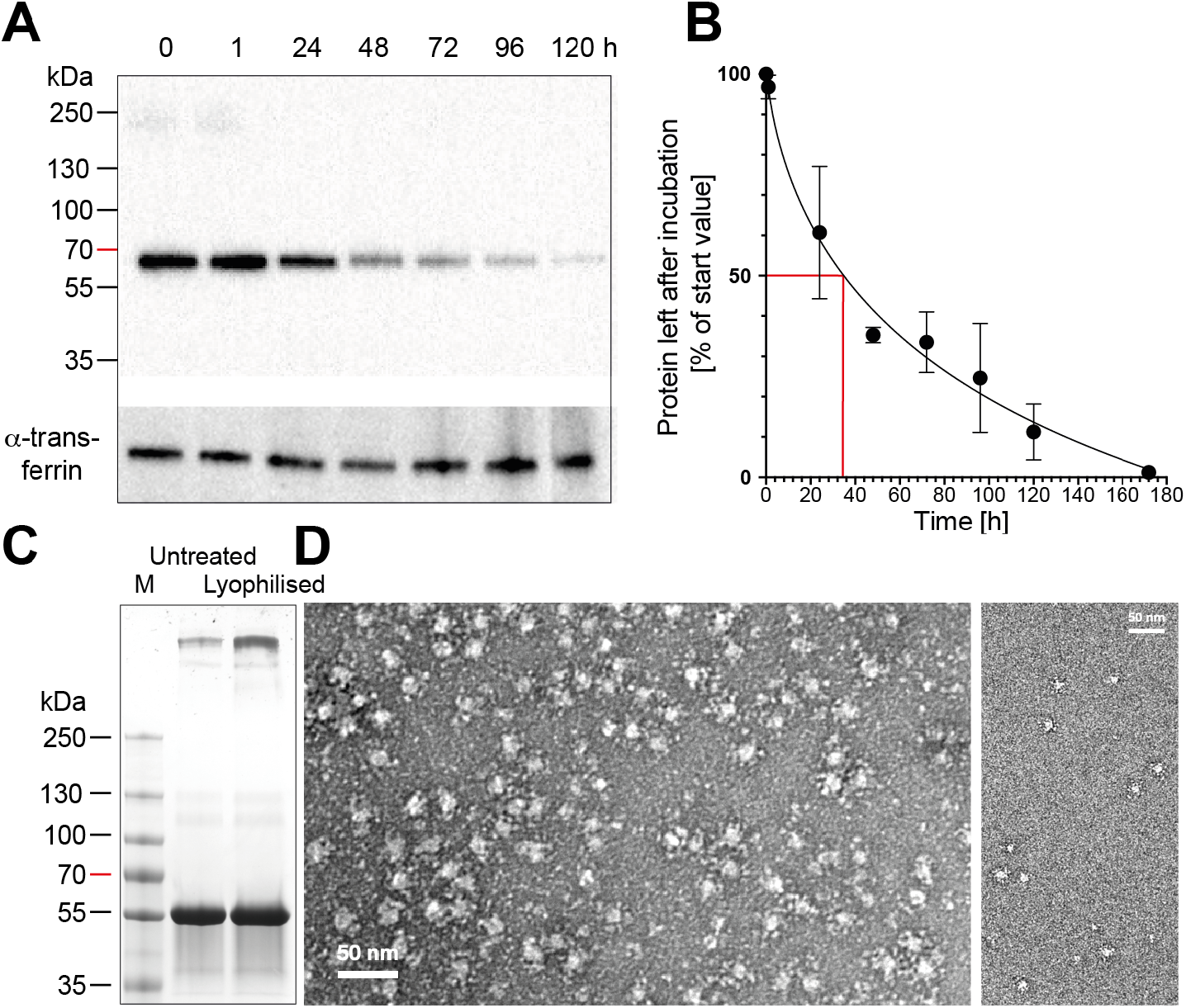
**A)** Plasma stability assay. RBD-S-Dps was incubated with non-heated human blood plasma for the amount of time indicated. SDS-PAGE and Western blot against the histidine tag on the protein. Transferrin was used as loading control and also detected by Western blot. **B)** Quantification of the data in A). The red line indicates the half-life of RBD-S-Dps under the conditions used. **C)** SDS-PAGE of RBD-S-Dps before and after lyophilisation. (Coomassie staining). **(D)** The lyophilised sample from C) was diluted to two different concentrations to demonstrate monodispersity and subjected to negative stain electron microscopy (left 20 x dilution and right 100 x dilution).

**Supplemental Figure 2.**
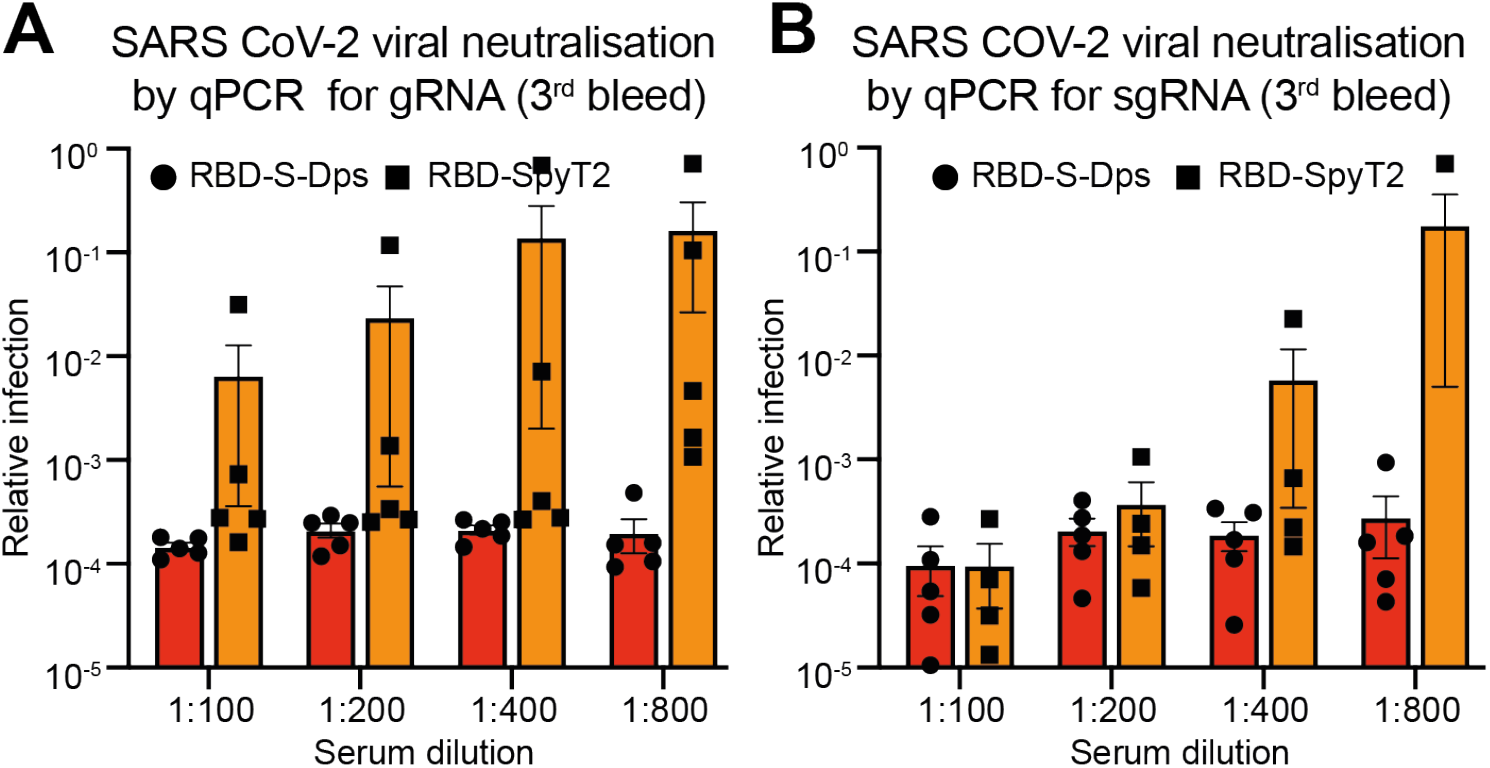
**A, B):** Vero cells expressing ACE2 and TMPRSS2 were infected with SARS-CoV-2 in the presence of serial dilutions of antisera. Viral replication was then determined after 24 h by RT-qPCR using probes for gRNA **(A)** or sgRNA **(B)**. Each point represents sera from an individual mouse.

**Supplemental Figure 3.**
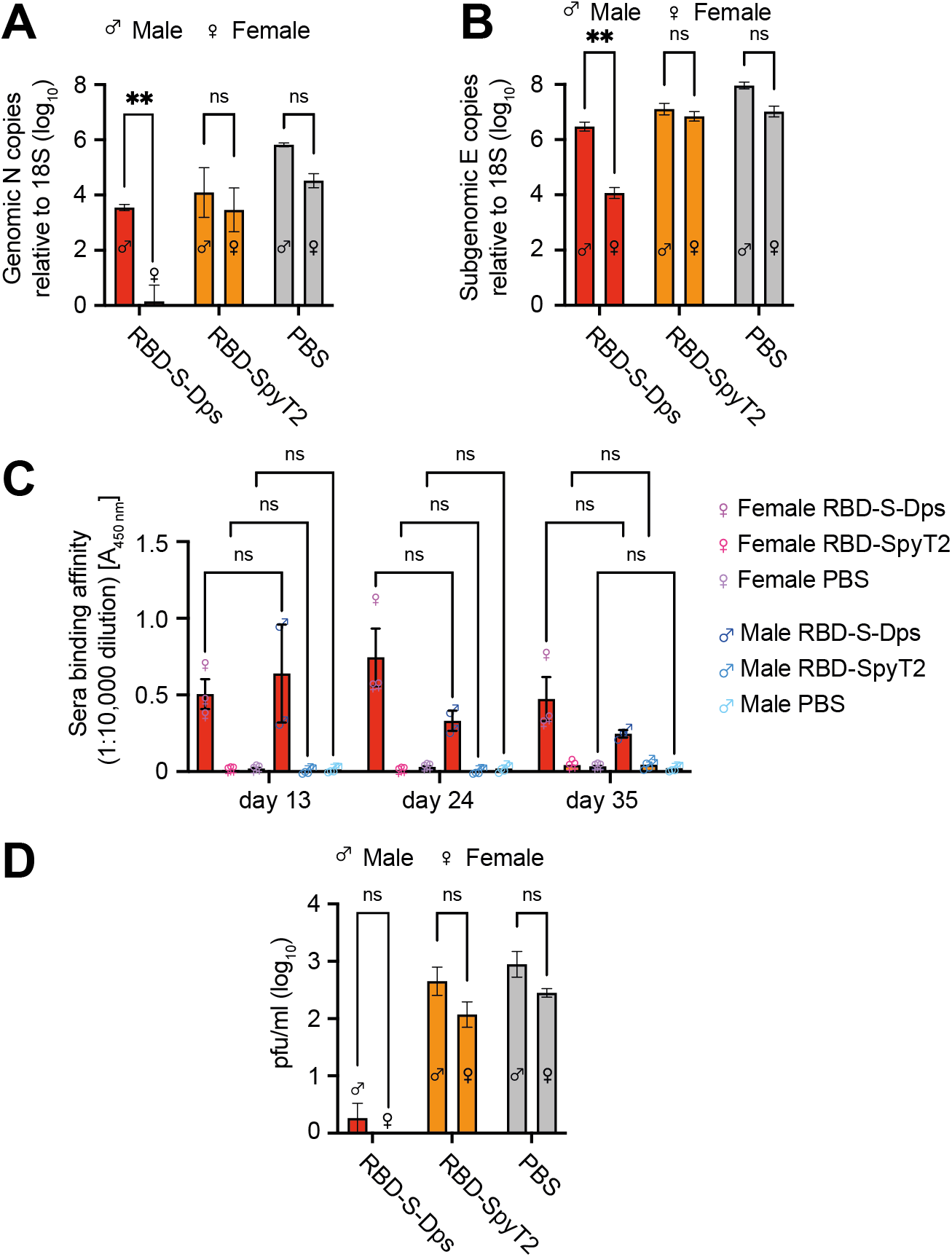
Mice were immunised with RBD-S-Dps, RBD-SpyT2 or given PBS control on day 1 and then challenged with SARS-CoV-2 on day 28. **A & B)** Genomic and subgenomic (gRNA, sgRNA) qPCR on RNA extracted from lung homogenates, using probes against NP or E, respectively. **C)** Sera from days 13, 24 and 35 were tested for anti-RBD antibodies by ELISA. Two-way ANOVA tests show that there are non-significant differences between male and female antibody responses. **D)** Plaque assay using lung homogenates from mice culled seven days post-infection.

**Supplemental Figure 4.**
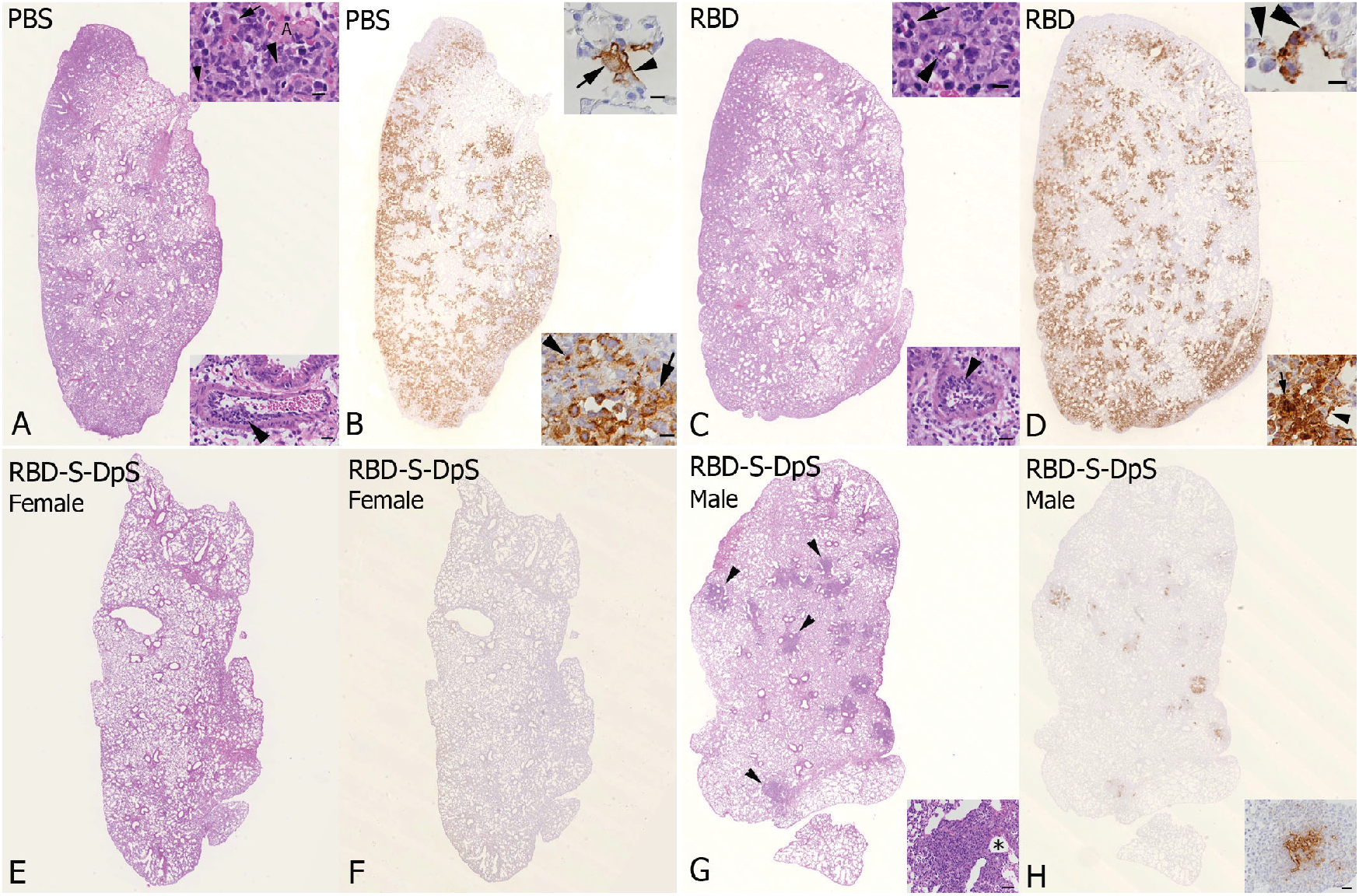
Lung, left lobe, K18-hACE2 mice at day seven post infection, SARS-CoV-2 challenge experiment. Histological changes and SARS-CoV-2 antigen expression. **A, B: mice injected with PBS control. A)** Overview of the lung lobe, with multifocal extensive cell-rich consolidated areas. Inset top: consolidated area with activated type II pneumocyte (small arrow), syncytial cells (large arrowhead) and infiltrating neutrophil (small arrowhead); A – alveolus (bar = 10 µm). Inset bottom: artery with leukocyte infiltration of the wall (arrowhead; arteritis, bar = 20 µm). HE stain. **B)** Extensive SARS-CoV-2 antigen expression is seen in multifocal patchy areas within and close to consolidated areas, in pneumocytes and occasional macrophages. Inset top: alveolus with viral antigen expression in type I (arrowhead) and type II (arrow) pneumocyte. Inset bottom: consolidated area with viral antigen expression in macrophages (arrow) and degenerate cells (arrowhead). Immunohistology, haematoxylin counterstain, bars = 10 µm. **C, D: Monomeric RBD-SpyT2-immunised mice. C)** Overview of the lung lobe, with multifocal extensive cell-rich consolidated areas. Inset top: consolidated area with several neutrophils (arrowhead) and occasional necrotic cells (arrowhead, bar = 10 µm). Inset bottom: artery with leukocyte infiltration of the wall (arrowhead; arteritis) and mild periarterial edema (bar = 20 µm). HE stain. **D)** Extensive SARS-CoV-2 antigen expression in multifocal patchy areas within and close to consolidated areas, in pneumocytes and occasional macrophages. Inset top: alveolus with viral antigen expression in pneumocytes of which some are degenerate (arrowheads). Inset bottom: consolidated area with viral antigen expression in macrophages (arrow) and type I pneumocyte (arrowhead). Immunohistology, haematoxylin counterstain; bars = 10 µm. **E-H: Multimerised RBD-S-Dps immunised mice**. **E, F.** Female animal. **E)** Overview of the lung lobe. The histological changes are restricted to focal areas of mildly increased interstitial cellularity. HE stain. **F**) There is no evidence of viral antigen expression. Immunohistology, haematoxylin counterstain. **G, H.** Male animal. **G)** Overview of the lung lobe, with several small, randomly distributed peribronchiolar cell-rich areas (arrowheads). Inset: Closer view of focal cell rich area (asterisk: bronchiole); bar = 50 µm, HE stain. **H**) SARS-CoV-2 antigen expression is restricted to the cell rich focal areas, in pneumocytes and occasional macrophages (inset). Immunohistology, haematoxylin counterstain; bar = 20 µm.

**Supplemental Figure 5.**
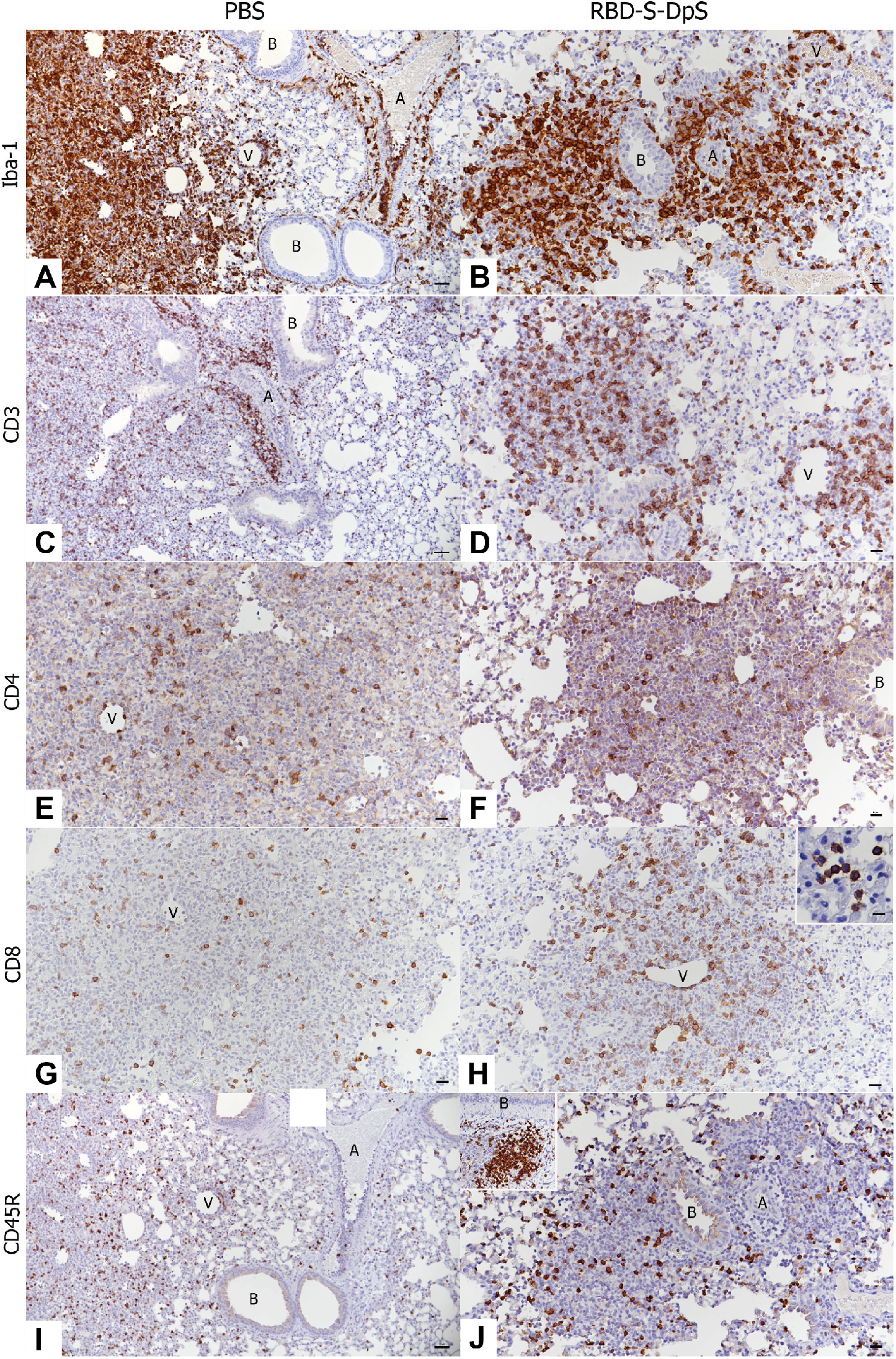
Lung, K18-hACE2 mice. Composition of the inflammatory infiltrates. **A, B:** staining for macrophages (Iba1+). **A)** PBS-control animal. Macrophages are the dominant infiltrating cells in the consolidated areas and in the vasculitis. A – artery with infiltration of the wall. V – vein with infiltration of the wall. B – bronchiole. Bar = 50 µm. **B)** RBD-S-Dps animal, male. Macrophages are the dominant infiltrating cells in the focal infiltrates. A – artery. V – vein. B – bronchiole. Bar = 20 µm. **C, D:** staining for T cells (CD3+). **C)** PBS-control animal. T cells are numerous in the consolidated areas and in the vasculitis. A – artery with infiltration of the wall and perivascular T cell accumulation. B – bronchiole. Bar = 50 µm. **D)** RBD-S-Dps animal, male. T cells are numerous in the focal infiltrates. V – vein. Bar = 20 µm. **E, F:** staining for CD4. **E)** PBS-control animal. Within the infiltrates, CD4 positive cells are numerous. V – vein. Bar = 20 µm. **F)** RBD-S-Dps animal, male. Within the infiltrates, CD4 positive cells are present in moderate number. B – bronchiole. Bar = 20 µm. **G, H:** staining for CD8. **G)** PBS-control animal. CD8 positive cells are less numerous. V – vein. Bar = 20 µm. **H)** RBD-S-Dps animal, male. CD8 positive cells are more abundant. V – vein. Bar = 20 µm. Inset: CD8 positive cells are also present in the lumen of several alveoli. Bar = 10 µm. **I, J:** staining for B cells (CD45R/B220+). **I)** PBS-control animal. B cells are observed in moderate numbers in the consolidated areas and are rare in the vasculitis. A – artery with infiltration of the wall. V – vein with infiltration of the wall. B – bronchiole. Bar = 50 µm. **J)** RBD-S-Dps animal, male. B cells (CD45R/B220+) are observed in moderate numbers in the focal infiltrates. A – artery. B – bronchiole. Bar = 20 µm. Inset: Focal peribronchial (B) B cell aggregate. Bar = 20 µm. Immunohistology, haematoxylin counterstain.

**Supplemental Table 1.**
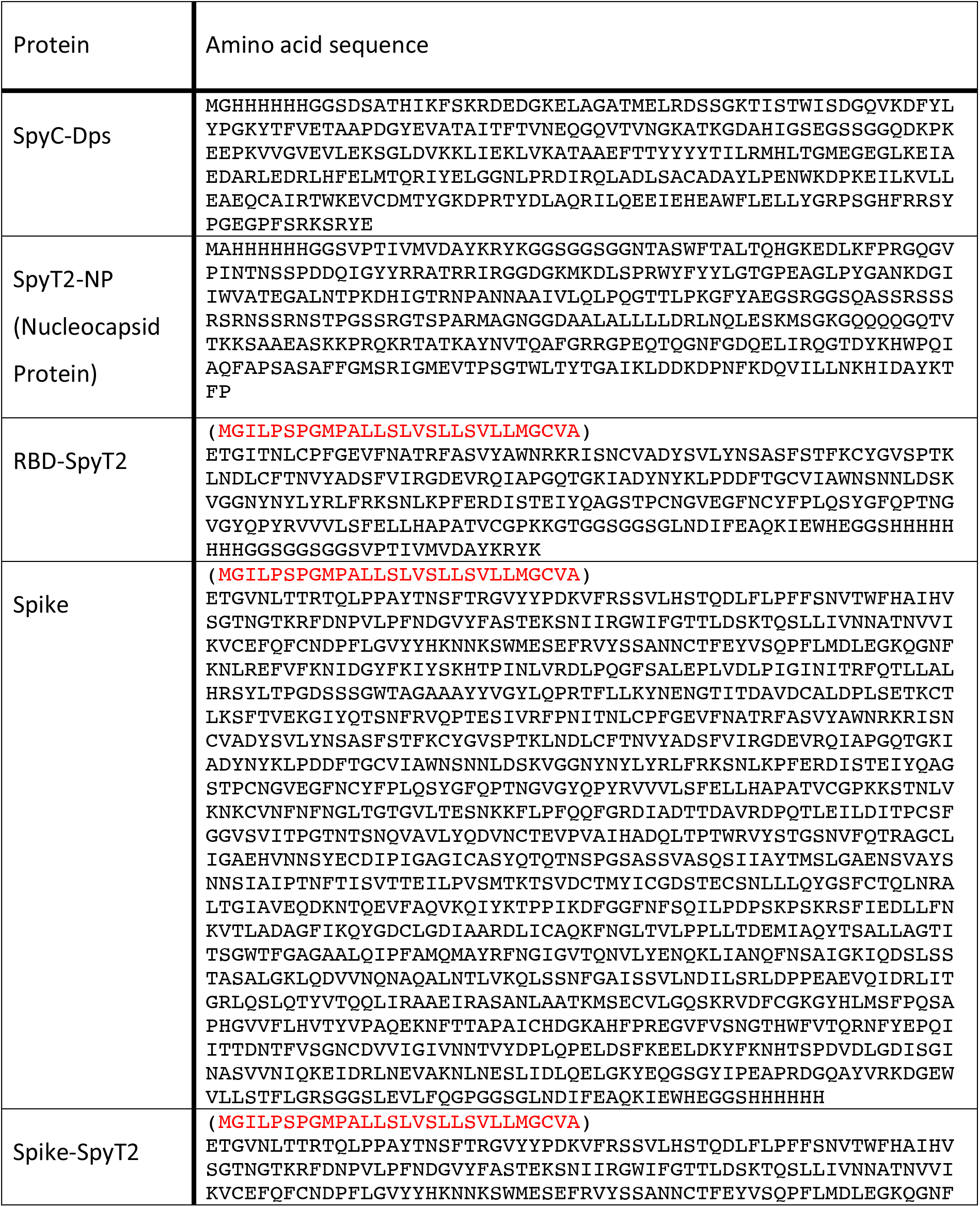
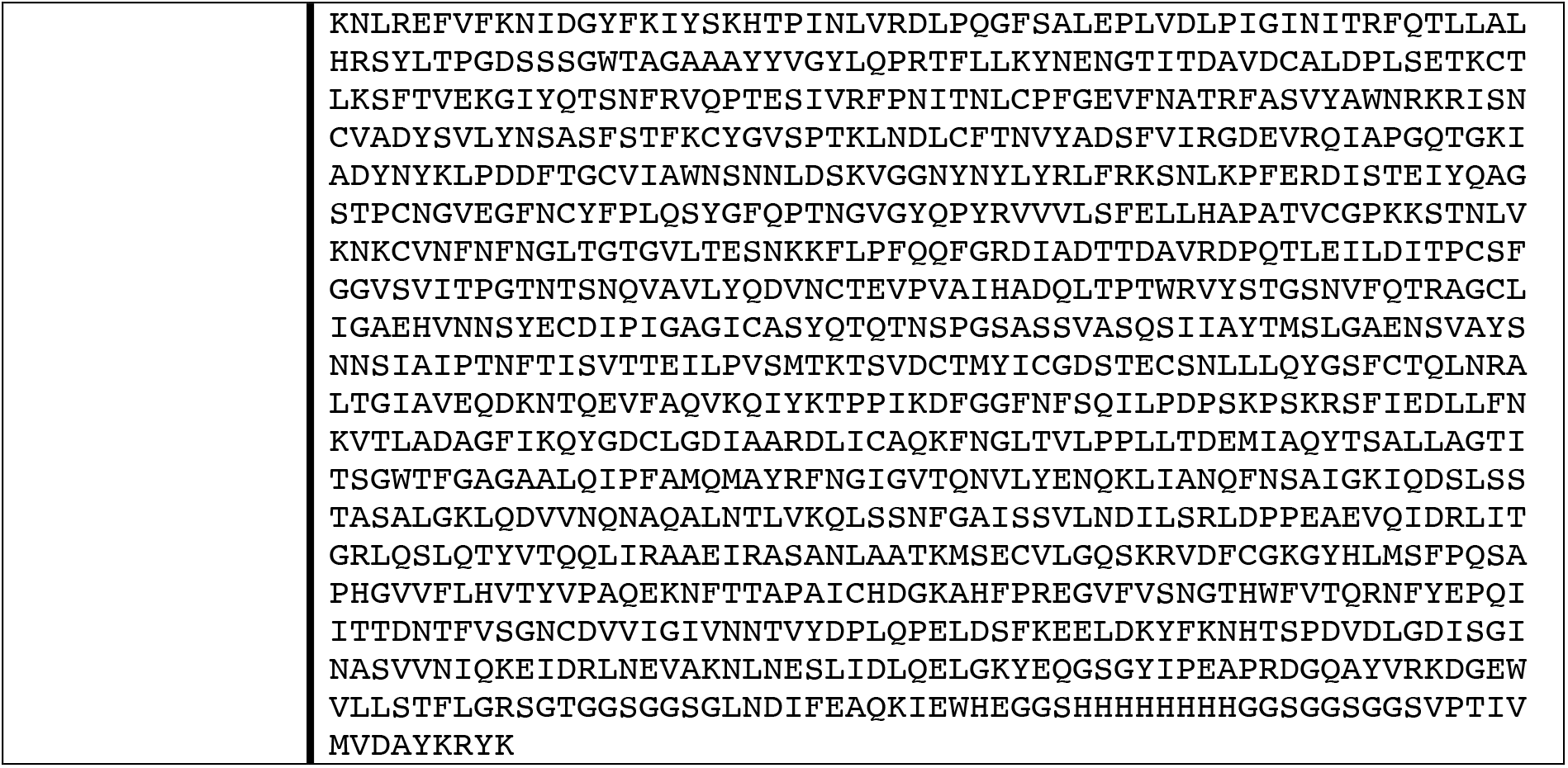
Amino acid sequences of the proteins used in this work. Signal sequences for secretion in mammalian cells are indicated in red.

